# A novel mechanism for volitional locomotion in larval zebrafish

**DOI:** 10.1101/189191

**Authors:** David E. Ehrlich, David Schoppik

## Abstract

To locomote stably, animals must coordinate volitional actions that change posture with opposing reflexes that hold posture constant [1–8]. These conflicting actions are thought to necessitate integrated control, in which reflexes are modulated to permit or even produce volitional movements [9–14]. Here we report that larval zebrafish (*Danio rerio*) utilize a simpler control scheme featuring independent volitional and reflexive movements. We present behavioral evidence that larvae swim in depth by appending destabilizing trunk rotations to steer with independent rotations to balance. When we manipulated buoyancy to deflect fish up or down, they redirected steering without coordinated changes to their balance reflex. As balance developed and increasingly opposed destabilization-mediated steering, larvae acquired compensatory use of their pectoral fins to steer. Removing the pectoral fins from older larvae impaired steering but preserved the strong balance reflex. Consequentially, older larvae without fins were strikingly less maneuverable — unable to revert to destabilization-mediated steering — revealing a rigidity inherent within the framework of independent volitional and reflexive control. Larval zebrafish therefore produce effective but inflexible locomotion by sequencing independent volitional and reflexive movements. These results reveal a simple control scheme, applicable for robotic design, that solves the general problem of coordinating volitional movements with the vital reflexes that oppose them.

---

Animals destabilize their bodies to move, complicating the challenge of maintaining balance [15]. For example, humans walk in “controlled falls,” toppling forward with each step before regaining stability [16]. The conflict between movement and stability is pervasive, but a simple body plan constrains the problem for young zebrafish. During the first days of swimming, zebrafish larvae possess largely ineffectual fins and simply propel where they point [17, 18]. Larvae must therefore rotate away from horizontal to climb or dive for a number of vital behaviors, including prey capture, predator evasion, buoyancy control, and circadian migration [19–21]. However, larvae also exhibit a righting reflex, a tendency to reorient towards a preferred posture near horizontal [22, 23] (Figure 1A). Larvae are therefore a simpler model system faced with a universal challenge: to integrate reflexes into locomotion so as to permit mobility but maintain stability.

**Figure 1:**
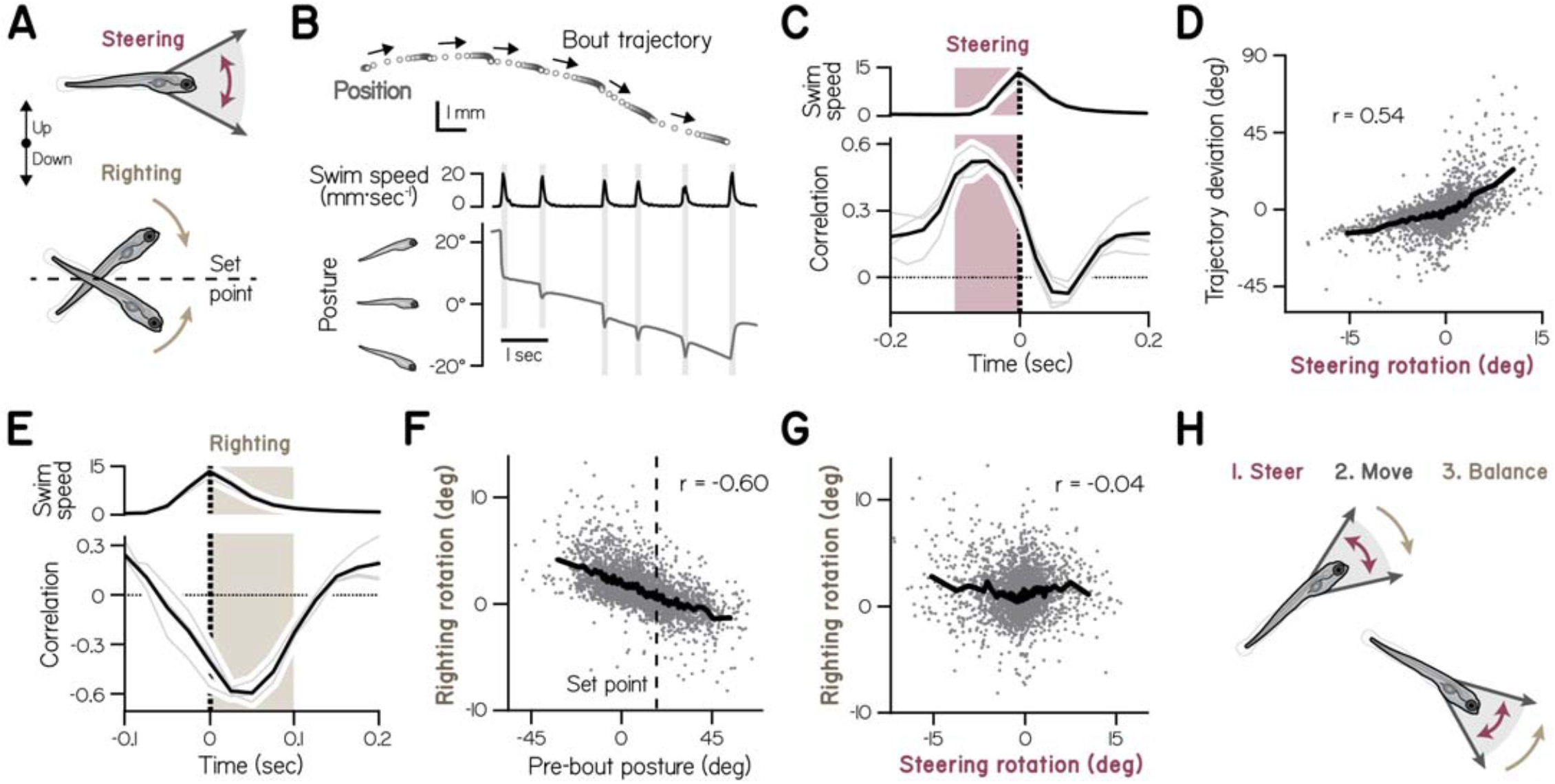
Zebrafish larvae swim with independent rotations that steer and balance. (A) Larvae propel where they point, and must steer in depth by rotating away from horizontal. An opposing righting reflex maintains them near horizontal, via rotations towards a preferred posture –the reflex “set point. (B) Positionsofa1 wpf larva during a representative swimming epoch. Trajectories of6swim bouts are depicted as arrows (top). Peaks of swim speed corresponding to the 6 bouts coincide with fast rotations of the body in the nose-up/down (pitch) axis (bottom). (C) The relationship (Pearson’s correlation coefficient) between instantaneous trunk rotations throughout the bouts with deviations from null trajectories. Bouts were aligned by peak swim speed (top). The red “steering window contains the largest positive correlation between changes to trajectory and posture. Data are plotted for individual clutches (sibling groups, thin lines, n=4) and their mean (thick line). (D) Deviation from the null trajectory is plotted versus trunk rotation across the steering window (red in C), for each of 3556 observed bouts. The line conveys means for bins of 50 bouts. (E) The relationship (Pearson’s correlation coefficient) between instantaneous trunk rotations with the pre-bout posture (100 msec before peak swim speed) for bouts aligned by peak swim speed (top). The tan “righting window contains the largest negative correlation. (F) Trunk rotation across the righting window (tan in E) is plotted as a function of pre-bout posture for bouts in (D). The empirical posture set point, computed as the best-fit line intercept, is indicated with a vertical dashed line. (G) Balancing rotations are plotted as a function of steering rotations for the bouts in (D). (H) Schematic depicting how larvae swim in depth – by first making steering rotations independent from initial posture, then attaining speed, and finally with righting rotations that are independent from how they steered.

Due to their small size, larvae swim intermittently, alternating between passive periods when they are rotated and pushed by external forces, and active bouts when they swim and adjust their posture [23–25]. Accordingly, freely-swimming larvae spontaneously vary their posture and depth, but tend to remain near horizontal (Figure 1B) [23]. Therefore, to define how larvae controlled their bodies to achieve both mobility and stability, we examined their posture during spontaneous swim bouts at 1 week post-fertilization (wpf). First, we examined when larvae made spontaneous trunk rotations that permitted changes to depth – volitional control of posture we termed “steering.” Second, we determined when larvae exhibited their righting reflex –responsesto instabilityin which they reoriented towards their preferred posture.

We found that larvae changed their elevation by steering up or down while they accelerated during bouts. Specifically, we examined whether rotations during a bout impacted its trajectory, or if larvae propelled where they pointed when a bout began (the null trajectory). Deviations from the null trajectory were positively correlated with trunk rotations during bouts, particularly those during the 100 msec before speed peaked (r=0.54; Figure 1C, 1D). This window corresponds to the greatest acceleration each bout (Supplemental Figure S1) and includes rotations that precede trunk rotation (Supplemental Figure S2). Therefore, larvae steer by rotating their trunks at the start of a bout.

Larvae temporally offset steering and their righting reflex, rotating towards their preferred posture while decelerating. In control theoretic terms, the reflex provides negative feedback to reduce postural error, the extent posture deviates from its “set point.” Postural error at the start of a bout was negatively correlated with rotations during the bout, indicating that bouts acutely stabilize posture (Figure 1E). In particular, the best correlated rotations occurred for 100 msec after speed peaked (r=-0.60; Figure 1F), corresponding to the greatest deceleration (Supplemental Figure S1). Larvae rotated towards an empirical set point, the posture at which no balance feedback was expected (intercept of the best-fit line to pre-bout posture and righting rotation), just nose-up to horizontal (17.4 ± 0.4°). Larvae therefore counteract destabilization, including volitional rotations away from horizontal, while they decelerate.

We tested whether the temporally dissociable rotations for steering and righting might be independent. Surprisingly, we found that steering and righting were not correlated (r=-0.04; Figure 1G). These data indicate that larvae locomote by sequencing nose-up/down rotations that steer with independent rotations that balance (Figure 1H). Accordingly, righting rotations did not correlate with trajectory changes (r=-0.08; Supplemental Figure S3A), suggesting larvae balance without directly impacting translation. Furthermore, steering rotations were not correlated with postural error before a bout (r=-0.02; Supplemental Figure S3B), such that movement is not constrained by stability. We therefore conclude that steering and righting are both sequential and independent.

We next tested whether steering and righting were under independent control. We measured how larvae maintain elevation when challenged - a problem that could be solved by modulating either steering or balance. To climb, for instance, a larva could steer upwards, away from its posture set point (Figure 2A, *lef);* alternatively, a larva could reorient nose-up using its righting reflex, by biasing the set point upwards (Figure 2A, *right)* [9, 26]. Complementarily, larvae could facilitate steering by weakening their reflex, permitting greater deviations from horizontal. Can larvae effectively control their swims while steering and righting independently?

**Figure 2:**
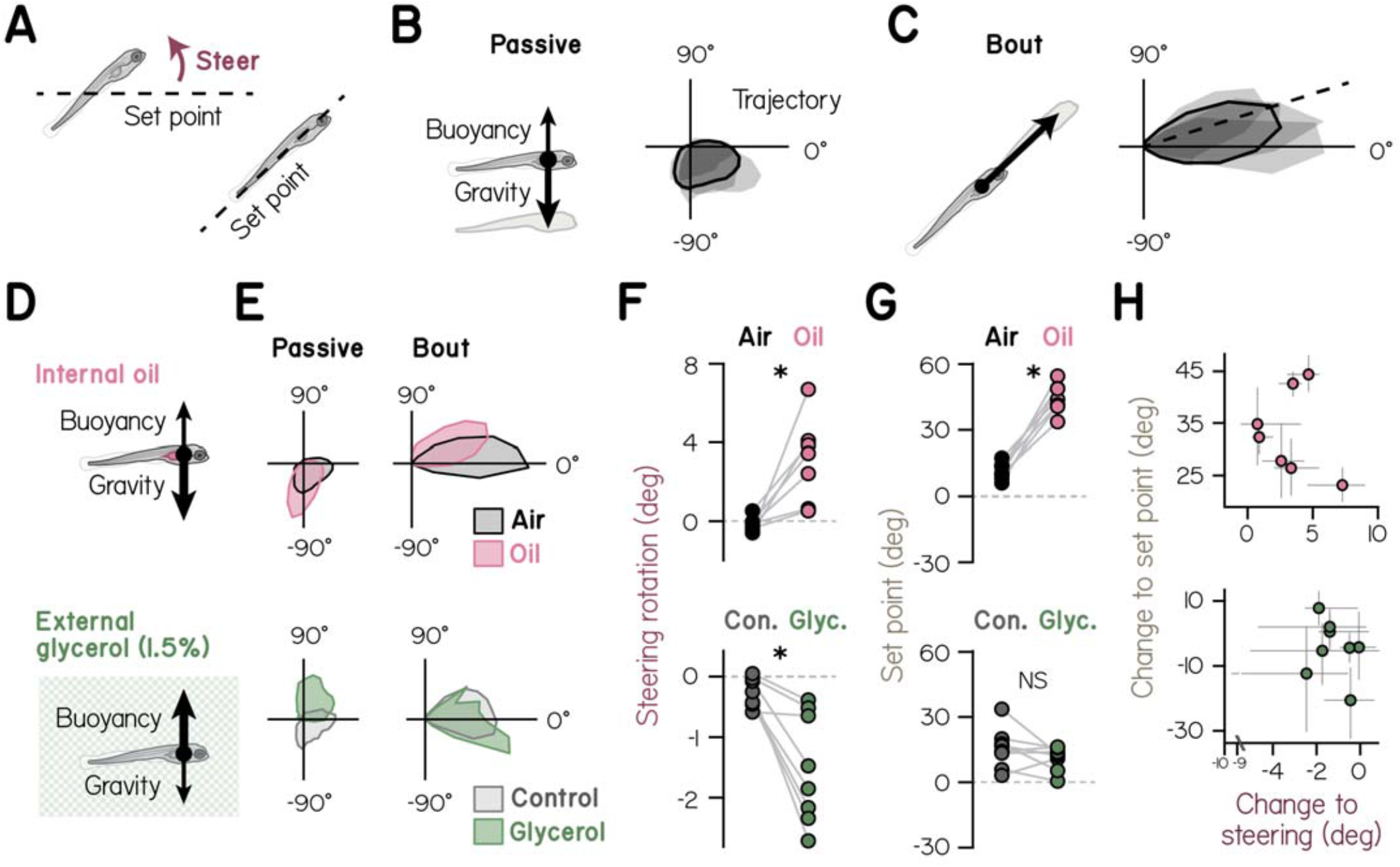
Larvae climb and dive by independently modulating their steering and righting rotations. (A) Larvae could theoretically climb by steering upwards away from their reflex set point (lef)oradjusting that set-point upwards (right). (B) Schematic of dominant forces – gravity and buoyancy– that act on larvae while passive between bouts. Polar probability distributions (right) show movement trajectories between bouts (at speeds<1 mm·sec-1) for individual clutches (shaded, n=4) and pooled data (line)at1 wpf. (C) Schematic of the dominant force – thrust – that acts on larvae during bouts. Polar probability distributions (right) show trajectories of bouts on identical scale to (B). The empirical reflex set point is indicated with a dashed line. (D) Larvae with oil-filled swim bladders were subjected to a greater force of gravity (top), while larvae swimming in glycerol were subjected to a greater force of buoyancy (bottom). (E) Polar histograms depict movement trajectories while passive (lef) and of bouts (right) for larvae with oil-filled swim bladders or siblings with air-filled swim bladders (top), as well as larvae in 1.5% glycerol and siblings in 0% glycerol (bottom). (F) Median steering rotations of larvae with oil-filled swim bladders or siblings with air-filled swim bladders (top, n=7 clutches of 8 larvae each), and larvae in 1.5% or 0% glycerol (bottom, n=8 clutches of 8 larvae each). * p¡0.025, paired t-test with Bonferroni correction. (G) Reflex set point for larvae with oil-filled swim bladders or siblings with air-filled swim bladders (top), as well as larvae in 1.5% glycerol and siblings in 0% glycerol (bottom). NS: not significant. (H) Effects of swim bladder (top) or glycerol manipulations (bottom) on reflex set point versus steering rotation, with bootstrapped 95% confidence intervals. Spearman’s rank correlation coefficients equal 0.11 (top) and 0.14 (bottom).

To challenge larvae to climb and dive, we leveraged their propensity to actively maintain position [27, 28]. During passive periods between bouts, larvae are moved by gravity and buoyancy (Supplemental Figure S4) [23]. They tend to be denser than water [29-31] and, consequently, tended to sink between bouts (Figure 2B) [23]. Under typical conditions larvae counteracted sinking by weakly biasing their bouts upwards, consistent with their natural nose-up bias to posture (Figure 2C).

If we displaced larvae vertically by manipulating their buoyancy, they biased swims to climb or dive. To exacerbate sinking we altered the swim bladder, a sac larvae typically inflate with air from the surface **∼**3 days post-fertilization [19]. Larvae instead inflate their swim bladders with denser paraffin oil if raised with oil on the water’s surface [23] (Figure 2D, *top)*. Larvae with oil-filled swim bladders sank more between bouts and biased those bouts systematically upwards, compared to siblings with air-filled swim bladders (Figure 2E, *top)*. In contrast, when we increased the force of buoyancy by placing larvae acutely in a low concentration (1.5%) of glycerol, larvae tended to rise between bouts and bias those bouts downwards (Figure 2D, 2E, *bottom)*.

Larvae could maintain elevation when challenged by modulating either their steering and/or their righting reflex. Denser larvae climbed by steering upwards (Figure 2F; paired t-test with Bonferroni correction: t_6_=3.87, p<0.025), while more buoyant larvae dove by steering downwards (t_7_=-4.22, p<0.025). In addition, denser larvae modulated their posture set point, using their righting reflex to skew posture, and therefore trajectory, upwards (p<0.025, t_6_=10.81; Figure 2G, top). However, larvae with elevated buoyancy did not adjust their set point (p>0.025, t_7_=-1.47; Figure 2G, *bottom)*, meaning set point modulation was selective.

If steering and righting are independent, larvae ought bias the former up or down without coordinating the reflex set point. Crucially, we found that individual changes to steering and set point were uncorrelated, whether larvae were climbing (Figure 2H; Spearman’s *ρ* = 0.11) or diving (*ρ* = 0.14). Furthermore, larvae did not aid steering by suppressing reflex strength, as measured by its gain - the proportion of postural error canceled by the average bout (Table S1; Oil vs. Air: t_6_=-1.78, p>0.025; 1.5% vs. 0% glycerol: t_7_=0.87, p*>*0.025). Together, these data show that when challenged, zebrafish larvae control elevation by independently modulating the volitional and reflexive components of locomotion.

Independent control of steering and righting poses a potential problem as fish mature. Young larvae make destabilizing rotations to steer, meaning developmental improvements to stability [23] will impair steering unless larvae compensate. However, if steering and righting mature independently, larvae would be unable to preserve mobility by ensuring that balance does not come to dominate. Evidence that larvae are unable to check their righting reflex to maintain mobility would support the hypothesis of independent control. Therefore, we investigated and manipulated how steering and righting change throughout early development.

We found that improvements to balance permit greater stability as fish develop. We measured locomotion in siblings from 1 to 3 wpf (n=4 groups of 8 fish), and found that older larvae prolonged the balance phase of their bouts (Figure 3A). With age, each bout came to reduce a greater proportion of postural error, equating to larger reflex gain (Figure 3B; main effect of age: F_(1,7)_=13.91, p<0.01; main effect of clutch: F(3,7)=2.66, p>0.05). When pooling all bouts, the gain doubled from 0.065 at 1 wpf (3556 bouts) to 0.129 at 3 wpf (709 bouts). Reflex gain was inversely correlated with posture variation, such that older larvae better constrained their trunks about their preferred posture (Figure 3C; Spearman’s *ρ* = -0.62, p<0.05). Maturation of the righting reflex therefore affords greater stability while swimming.

**Figure 3:**
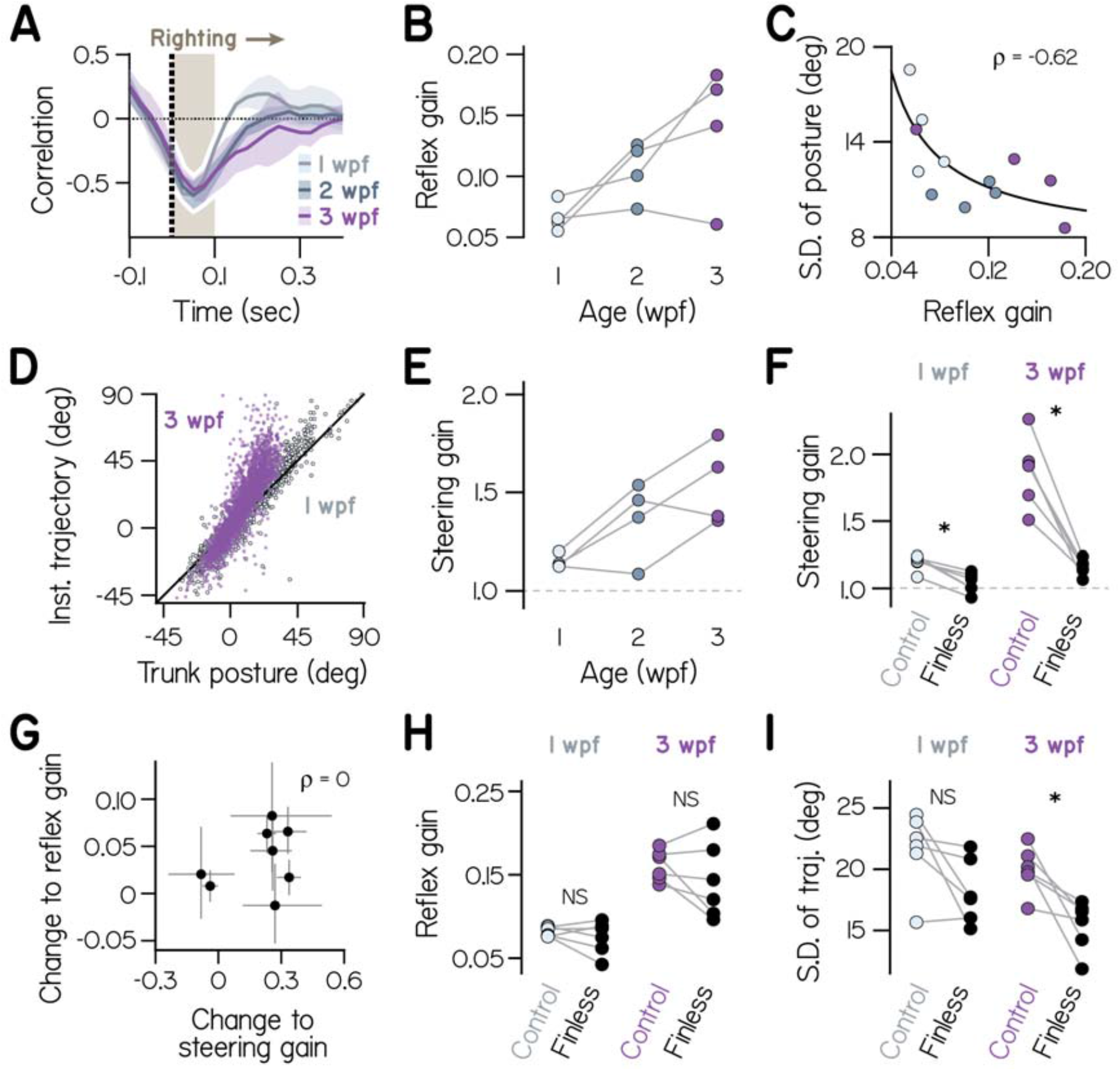
Steering and righting mature independently. (A)As inFigure 1E, Pearson’s correlation coefficient relates instantaneous trunk rotations with the pre-bout posture. Negative correlations are prolonged with age, plotted as the mean and S.D. across 4 clutches at each of 1, 2, and 3 weeks post-fertilization (wpf). (B) Reflex gain is plotted for 4 clutches measured at 1, 2, and 3 wpf. (C) Standard deviation of posture is plotted as a function of reflex gain, with a least-squares fit inverse function and Spearman’s rank correlation coefficient listed. (D) Trajectoryatpeak speed of each boutisplottedas a function of concurrent trunk posture for bouts at 1wpf (light blue) and 3 wpf (dark blue), with the unity line in black. (E) Steering gain, the Theil-Sen slope estimate for data in (D), is plotted for 4 clutches at 1, 2, and 3 wpf. (F) Steering gain is plotted for larvae with surgically removed pectoral fins and unaltered siblings (n=6) at 1 and 3 wpf. Two-way ANOVA results in text; * p¡0.025, paired t-test with Bonferroni correction. (G) Changes to reflex gain and steering gain from 1-2 or 2-3 wpf across the 4 clutches, with bootstrapped 95% confidence intervals and Spearman’s rank correlation coefficient listed. (H) Reflex gain for larvae with surgically removed fins and unaltered siblings at 1 and 3 wpf. Two-way ANOVA results in text; NS: not significant, p>0.025. (I) Standard deviation of observed bout trajectories for larvae with surgically removed fins and unaltered siblings at 1 and 3 wpf. Two-way ANOVA results in text.

As stability increased, larvae maintained maneuverability through parallel improvements to steering. At one wpf, larvae propelled where they pointed, such that steering rotations provided a proportional change to swim trajectory - corresponding to steering gains near 1 (Figure 3D,3E; Theil-Sen estimated slope of trajectory vs. posture; Supplemental Figure S5). Older larvae could effectively climb or dive while remaining closer to horizontal, having gained additional control over their movements. They did not simply propel where they pointed, but skewed trajectories to amplify steering rotations, giving a steering gain much larger than 1 (Figure 3D,3E; two-way ANOVA, main effect of age: F(1,7)=22.73, p<0.01; main effect of clutch: F(3,7)=3.95, p>0.05). Therefore, developing larvae came to swim with less trunk destabilization, much as human toddlers walk more stably by learning to swing their arms [32], and gymnasts learn to prioritize trunk stability in their movements [33].

We found that steering improvements reflect emergent fin use. Mature fish use their pectoral fins, homologues of amniote forelimbs, to steer in depth [29, 34-36], but week-old zebrafish possess minimally functional pectoral fins [18]. After surgical removal of the pectoral fins, all larvae propelled precisely where they pointed, with steering gains reduced to 1 (Figure 3F; two-way ANOVA, main effect of fin removal: F_(1,20)_=57.37, p<0.001; main effect of age: F(1,20)=49.81, p<0.001; interaction effect: F(1,20) =24.78, p<0.001; n=6 clutches). Fin removal also abolished the acquired agility to make large trajectory changes from one bout to the next (Supplemental Figure S6). While the effect of fin removal was large at 3 wpf, cutting steering gain nearly in half (paired t-test with Bonferroni correction, t5=730, p<0.001), larvae at 1 wpf also exhibited significant impairment (t5=5.22, p<0.001). For fin use to increase steering gain, larvae must skew movements upwards when their trunks are oriented up, and downwards when oriented down, suggesting larvae actively coordinate their trunks and fins to steer. Together, these data suggest larvae increasingly use pectoral fins to amplify the effects of trunk rotation on their swimming movements.

We found that fin development selectively impacted steering. Aside from effects on the direction of locomotion, larvae swam comparably after loss of the fins. Fin removal had no impact on their rate of swimming or the speed and displacement of their bouts (Supplemental Table S2). Furthermore, larvae produced comparable trunk rotations after loss of the fins (Supplemental Table S2; main effect of fin removal on angular speed: F(1,20)=0.13, p>0.05). The pectoral fins therefore complement trunk-based steering but are not necessary to rotate the trunk. Importantly, fin-derived improvements to steering progressed distinctly from the maturation of the righting reflex, as changes to steering and reflex gains were uncorrelated throughout development (Figure 3G; Spearman’s *ρ* = 0). Across clutches, larvae improved steering and righting to different extents and at different rates (Figure 3B,3E). Thus steering and righting develop complementarily, even as their control remains independent.

Independent control made larvae inflexible, unable to adapt steering when challenged. Specifically, older larvae could not modulate their reflex to restore effective trunk-based steering when their fins were removed. Forced once more to propel where they point, finless larvae at 3 wpf must deviate from horizontal to climb or dive. However, they failed to reduce their reflex gains after fin removal (Figure 3H, Supplemental Figure S7A; main effect of fin removal: F_(1,20)_=1.36, p*>*0.05; main effect of age: F_(1,20)_=46.10, p<0.001) and therefore continued to tightly restrict trunk posture after fin removal (Supplemental Figure S7B; two-way ANOVA of posture variance by clutch, main effect of age: F(1,20)=40.34, p<0.001; main effect of fin removal: F(1,20)=0.23, p>0.05). Greater stability at 3 wpf therefore equated to less maneuverability without the fins, evident in reduced variation of swim trajectory (Figure 3I; main effect of fin removal: F_(1,20)_=15.47, p<0.001; paired t-test with Bonferroni correction at 3 wpf: t5=3.6, p<0.025; at 1 wpf: t5=2.3, p>0.05). By reverting larvae to undeveloped steering, we discovered a limitation of their locomotion – age-expected volitional and reflexive control are necessary for proper swimming, as the two are age-matched to achieve stability and maneuverability.

By producing effective locomotion across a range of parameters for steering and righting, independent control allowed larvae to modify existing movement patterns, a prerequisite for locomotor development [37–39]. However, independent control left larvae unable to manage the trade-off between stability and maneuverability, to compensate for fin loss by reverting to the trunk-based steering of younger larvae. Temporal dissociation thus provides a simple but limited means for new swimmers to avoid conflict between steering and stabilizing movements.

Temporary suppression of righting reflexes may be a useful and conserved mechanism to sequence volitional and reflexive control. Cycles of de- and re-stabilization are also apparent in human gait [16]. Vestibular stimulation impacts stepping late but not early in the gait cycle [40], when reflexes may be suppressed to avoid interference with volitional control of walking. A neural means to inactivate vestibular reflexes has been described during voluntary movements of the eyes [41–45], and an analogous circuit may exist to inactivate descending vestibular commands during locomotion. Modulation of sensory signals according to gait cycle phase is a general mechanism to appropriately adjust locomotion to reflect ongoing sensation [46].

In summary, our experiments show that effective locomotion, in this case stable maneuvering under water, can emerge from an unexpectedly straightforward control scheme with independent volitional and reflexive movements. All animals that move themselves must permit instability in some form [9]; the larval zebrafish provides a natural proof-of-principle for one of the simplest possible solutions to this general problem – destabilize oneself to move, then re-stabilize when finished. Larvae also reveal one shortcoming of such rudimentary control, lacking the flexibility to revert to earlier modes of locomotion as they develop. Requiring no coordination beyond that which sequences volitional and reflexive components, independent control islikely simplerto neurally implement than the integrated control of more advanced locomotion [6, 7], and thus well-suited for robotic control. Animals that move and then balance may be a developmental and evolutionary intermediary between those that move without balance and those that move and balance in concert.

## Methods

### Fish Care

All procedures involving zebrafish larvae (*Danio rerio*) were approved by the Institutional Animal Care and Use Committee of New York University. Fertilized eggs were collected from in-crosses of a breeding population of Schoppik lab wild-type zebrafish maintained at 28.5°C on a standard 14/10 hour light/dark cycle. Before 5 dpf, larvae were maintained at densities of 20-50 larvae per petri dish of 10 cm diameter, filled with 25-40 mLE3with 0.5 ppm methylene blue. Subsequently, larvae were maintained on system water in 2-L tanks at densities of 6-10 per tank and fed twice daily. Larvae received powdered food (Otohime A, Reed Mariculture, Campbell, CA) until 13 dpf and brine shrimp thereafter. Larvae were checked visually for swim bladder inflation before all behavioral measurements.

### Physical manipulations

To generate larvae with swim bladders filled with paraffin oil, 3 dpf larvae were visually checked for the absence ofswim bladders and transferred to 50 mL conical tubes (Falcon, Thermo Fisher Scientific) at a density of 12 larvae per tube, as previously [23]. The tubes were filled with 45 mL E3 containing methylene blue (as above), then topped with 5 mL paraffin oil (VWR, Radnor, PA) and incubated until 5dpf for experimentation. Control siblings were maintained similarly in 50 mL E3 in conical tubes without an oil surface.

Pectoral fins were removed surgically from larvae anesthetized in 0.02% ethyl-3-aminobenzoic acid ethyl ester (MESAB, Sigma-Aldrich E10521, St. Louis, MO). Pairs of anesthetized, length-matched siblings were immobilized dorsal-up in 2% low-melting temperature agar (Thermo Fisher Scientific 16520), and both pectoral fins of one larva were removed by pulling the base laterally with forceps. Then, the altered larva and its control sibling were freed from the agar with a scalpel and allowed to recover in E3 for 4-5 hours prior to behavioral measurement.

### Swimming measurement

Data were analyzed from a previous study of 8 larvae from each of 4 clutches at 1 wpf, and from larvae with oil-filled swim bladders and control siblings at 5 days post-fertilization [23]. For measuring effects of pectoral fin removal, new data were captured identically from 6-8 larvae per condition per clutch (n=7) at 1 and 3 wpf. For comparing swimming in 0 and 1.5% glycerol solutions, 8 larvae per condition per clutch (n=8). Siblings were transferred to a glass tank (93/G/10 55x55x10 mm, Starna Cells, Inc., Atascadero, CA) filled with 24-26 mL E3 and recorded for 24 hours unless otherwise noted. The thin tank (10 mm) maximized the time the fish spent swimming in the imaging plane. The enclosure containing the tank was kept on the same 14/10 hour light/dark cycle as the aquaculture facility using overhead LEDs, which maintained water temperature at 26°C. Video was captured usingadigital camera (BFLY-PGE-23S6M, Point Grey Research, Richmond, BC, Canada) equipped withaclose-focusing, manual zoom lens (18-108 mm Macro Zoom 7000 Lens, Navitar, Inc., Rochester, NY, USA) with f-stop set to 16 to maximize depth of focus. The field-of-view, approximately 2x2 cm, was aligned concentrically with the tank face. A5W940nm infrared LED backlight (eBay) was transmitted through an aspheric condenser lens with a diffuser (ACL5040-DG15-B, ThorLabs, NJ), and an infrared filter (43-953, Edmund Optics, NJ) was placed in the light path before the imaging lens.

Larvae with visually-confirmed paraffin oil-filled swimbladders and control siblings were tested in parallel as above, 8 larvae per clutch per condition (n=7), for 24 hours starting the day of 5 dpf. When imaging larvae with oil-filled swim bladders, E3 in the tanks was topped with a thin layer of paraffin oil, outside the field of view, to prevent supplementary inflation with air over the course of testing.

To increase the density of the swimming medium, larvae were acutely imaged in solutions of glycerol (Sigma-Aldrich, St. Louis, MO). Three sets of 8 siblings from each clutch (n=8 clutches) were measured in parallel at 7 dpf, in three concentrations of glycerol dissolved in E3 (0, 1.5, and 3% by volume). Data were collected for only 2 hours in glycerol solution to minimize physical buoyancy adaptation.

### Video acquisition and detection

Video acquisition was performed as previously [23]. Digital video was recordedat40Hzwithanexposure timeof1ms, and kinematic data were extracted online using the NI-IMAQ vision acquisition environment of LabVIEW (National Instruments Corporation, Austin, TX, USA). Background images were subtracted from live video, intensity thresholding and particle detection were applied, and age-specific exclusion criteria for particle maximum Feret diameter (the greatest distance between two parallel planes restricting the particle) were used to identify larvae in each image. The position of the visual center of mass and the trunk posture (orientation of the trunk in the pitch, or nose-up/down, axis) were collected each frame. Trunk posture was defined as the orientation, relative to horizontal, of the line passing through the visual centroid that minimizes the visual moment of inertia, such that a larva with trunk posture zero has its longitudinal axis approximately horizontal.

### Behavior analysis

Data analysis and modeling were performed using Matlab (MathWorks, Natick, MA, USA). Epochs of consecutively saved frames lasting at least 2.5 sec were incorporated in subsequent analyses if (1) they contained only one larva and (2) were captured during the last 12 hours of the light phase of the light-dark cycle, to minimize effects of light onset. Instantaneous differences of body particle centroid position were used to calculate swim speed. Consecutively detected bouts faster than 13_3_<u>^1^</u> Hz were merged into single bouts. Larvae with paraffin oil-filled swimbladders infrequently sank at speeds exceeding 5 mm/sec. For these larvae and control siblings with air-filled swimbladders, bouts were excluded if the movement vector was pointed vertically down (below -70° to horizontal; about 1% of bouts).

Bouts were excluded if initiated from a vertical orientation (*>*80°or <-80°, constituting 54 of 23,701 total bouts) due to ambiguous detection of vertical up and down postures. The posture set-point was calculated as the y-intercept of the best-fit line to pre-bout posture vs. late change in posture. The gain of the righting reflex was computed as the opposite of the slope of the best fit line of posture change (from the time of maximal bout speed to 250 msec later) to the pre-bout posture (100 msec prior to maximal speed). Developmental trajectories were analyzed by Two-way ANOVA treating clutch as a categorical factor and age as a continuous factor. Trajectory of a swim bout was defined as the direction of the translation vector from 100 msec before to 125 msec after max speed was reached. The change in trajectory was the difference between this trajectory and the pre-bout posture, such that a change in trajectory of zero described a larvae that swam directly where it pointed at the start of a bout. Changes in trajectory between successive bouts were compared based on fits to the mean cumulative probability distributions for clutches at a given age. Scale factors of the best-fit Weibull distributions were analyzed with two-way ANOVA and pairwise t-tests with Bonferroni correction.

### Data sharing

All raw data and analysis code are available online at http://www.schoppiklab.com/

## Acknowledgments

Research was supported by the National Institute on Deafness and Communication Disorders of the National Institutes of Health under award number DC012775. The authors would like to thank Katherine Harmon, Marie Greaney, Basak Sevinc, Eva Lancaster, Fernando Fuentes, Tim Gerson, Shane Hunt, and Belinda Sun for assistance with animal husbandry; Simon Sun for apparatus construction; Katherine Nagel, Michael Long, Rob Froemke, Nicolas Tritsch, Kishore Kuchibhotla, Katherine Eyring, Damon Lamb, Sam McKenzie, and members of the Schoppik and Nagel labs for helpful comments.

## Author Contributions

Conceptualization: DE and DS, Methodology: DE and DS, Investigation: DE, Visualization: DE, Writing: DE, Editing: DE and DS, Funding Acquisition: DS, Supervision: DS.

